# Branched-chain amino acid catabolism promotes ovarian cancer cell proliferation via phosphorylation of mTOR

**DOI:** 10.1101/2024.10.15.618560

**Authors:** Hannah J. Lusk, Monica A. Haughan, Tova M. Bergsten, Joanna E. Burdette, Laura M. Sanchez

## Abstract

Ovarian cancer is the sixth leading cause of cancer-related mortality among individuals with ovaries, and high-grade serous ovarian cancer (HGSOC) is the most common and lethal subtype. Characterized by a distinct and aggressive metastatic pattern, HGSOC can originate in the fallopian tube with the transformation of fallopian tube epithelial (FTE) cells, which metastasize to the ovary and subsequently to the omentum and peritoneal cavity. The omentum is a privileged metastatic site, and the metabolic exchange underlying omental metastasis could provide enzyme or receptor targets to block spread. In this study, we adapted a mass spectrometry imaging (MSI) protocol to investigate spatial location of 3D cocultures of tumorigenic FTE cells when grown in proximity to murine omental explants as a model of early metastatic colonization. Our analysis revealed several altered metabolites in tumorigenic FTE/omentum cocultures, namely changes in branched-chain amino acids (BCAA), including valine. We quantified the heightened consumption of valine, other BCAAs, and other amino acid-derived metabolites in omental cocultures using LC-MS assays. Our analysis revealed that metabolite concentrations when monitored with MSI from cell culture media in living culture systems have notable considerations for how MSI data may produce signatures that induce ionization suppression. Supplementation with valine enhanced proliferation and mTOR signaling in tumorigenic FTE cells, suggesting the potential of BCAA’s as a nutrient utilized by tumor cells during omental colonization and a possible target for metastasis.

**SIGNIFICANCE:** This study uncovers altered amino acid metabolism, specifically increased BCAA catabolism, at the interface of ovarian cancer cells and omental tissue in a coculture model of HGSOC secondary metastasis. Enhanced BCAA catabolism may promote cancer cell proliferation through mTOR signaling, presenting potential therapeutic value. These findings deepen our understanding of HGSOC pathogenesis and the metastatic tumor microenvironment, offering insights for developing new treatment strategies.

## INTRODUCTION

Ovarian cancer (OvC) is the most lethal gynecologic malignancy.(1) Projections indicate that in 2024, the United States will see 19,680 new cases and 12,740 deaths attributed to OvC.(1) Despite the term ‘ovarian cancer’ implying a singular condition, it encompasses several histologically distinct subtypes; among them, high-grade serous ovarian cancer (HGSOC) is the most prevalent and lethal.(2) HGSOC poses challenges in diagnosis due to non-specific symptoms and the absence of efficient screening methods.(2–4) Due to a lack of routine screening, approximately 80% of cases remain undetected until reaching metastatic stages, when the five-year survival rate drops below 30%.(2,3,5) This data underscores the need to investigate the molecular events driving HGSOC pathogenesis to facilitate the discovery of untapped therapeutic targets.

The metastatic behavior of HGSOC suggests that the omentum is a unique metastatic location, with omental metastases observed in approximately 80% of cases.(6,7) Several studies provide evidence for an “activated” phenotype of the peritoneal microenvironment associated with OvC, suggesting that chemical messengers specifically account for the preferential spread of fallopian tube derived HGSOC to the omental tissues during metastasis. The omentum is comprised of a variety of cell types, including adipocytes, fibroblasts, and macrophages, which can be dynamically converted to “cancer-associated” phenotypes, which is a process known to play a roles in disease progression.(8–10) For example, following co-culture with OvC cells, activated omental adipocytes have been shown to enhance the secretion of various adipokines, thus facilitating OvC dissemination.(7) A study by Rizi *et al*. demonstrated that cancer-associated adipocytes can also release arginine, which OvC cells may uptake, leading to increased nitric oxide production. Consequently, OvC cells release citrulline, a byproduct of nitric oxide synthase activity, which promotes adipogenesis in adipocytes.(11) Furthermore, cancer-associated adipocytes have been reported to transfer fatty acids to OvC cells, thereby promoting β-oxidation and energy generation which ultimately drives metastatic growth.(7) OvC cells also secrete soluble metabolites and microRNA that induce metabolic reprogramming in fibroblasts, prompting the acquisition of cancer-associated phenotypes. For instance, a study by Jeon *et al*. unveiled that lysophosphatidic acid, found in the malignant ascites of advanced-stage OvC patients, triggers the differentiation of adipose-derived mesenchymal stem cells into cancer-associated fibroblasts (CAFs).(12) Bidirectional interactions between OvC cells and CAFs have been reported in several studies.(13,14) In one such study, Wang *et al*. showed that CAFs confer platinum resistance to OvC cells by releasing glutathione and cysteine into the peritoneal microenvironment.(15) This finding is significant, as the standard ovarian cancer treatment includes surgical tumor removal followed by platinum-based chemotherapy.(16)

While we have some knowledge about the role secreted local factors play in omental colonization (14,17), the role of secreted metabolites, their spatial resolution, and their quantification has typically not been studied utilizing an untargeted label-free approach to detect the chemical cues driving this critical step in metastasis. Mass spectrometry imaging (MSI) is a useful tool for visualizing the spatial distribution of metabolites in biological samples.(18,19) Our lab previously developed an MSI protocol for analyzing 3D cocultures of explant murine tissues and mammalian cells and discovered and validated the presence of norepinephrine as a chemoattractant released from the ovary.(20,21) Further, our labs demonstrated that norepinephrine release was partially mediated by SPARC and impacted survival in a murine model.(22,23) In this present study, we adapted our MSI technique for use with omental tissue to investigate metabolic exchange in secondary metastasis.

MSI analysis of 3D omental cocultures led to the identification of several signals that appeared to be altered in tumorigenic conditions. MSI on cocultures in a divided chamber format revealed the site of origin of these signals, with some signals originating from tumorigenic FTE cells and others originating from omental tissue. Further analytical validation of *m/z* 118 revealed that it represented the branched-chain amino acid (BCAA) valine. Quantification of BCAAs and 21 other amino acids in coculture extracts demonstrated that valine and other BCAAs are actually consumed at higher levels in tumorigenic FTE/omentum cocultures compared to either condition alone or a non-tumorigenic fallopian tube cell model. Analysis of valine standard curves via MSI illustrated that the increase in signal observed was attributed to ion suppression, owing to the high concentration of valine in the media. Proliferation assays revealed that valine supplementation increases the proliferation of tumorigenic FTE cells, and immunoblotting showed that valine can stimulate the phosphorylation of mTOR. This work demonstrates the benefit of exploring spatial metabolic changes in cancer metastasis.

## MATERIALS AND METHODS

### 1. Mouse colony and omentum removal

All animals were treated in accordance with NIH Guidelines for the Care and Use of Laboratory Animals and the established Institutional Animal Use and Care protocol at the University of Illinois Chicago. Omental tissue was collected from female CD-1 mice 6-8 weeks in age (Charles River). Mice were housed in a temperature and light-controlled environment (12 hours light and 12 hours dark) and provided food and water *ad libitum*. Omentums were removed immediately after sacrifice and bisected using a dissecting microscope (Leica MZ6).

### 2. Cell lines

Dr. Barbara Vanderhyden from the University of Ottawa generously provided spontaneously immortalized murine ovarian surface epithelial (MOSE) cells and murine oviductal epithelial cells (MOE), which are equivalent to human FTE. These MOE cells were modified in our lab to express different genetic variations, including a scrambled control shRNA (MOE SCR^shRNA^) and an shRNA targeting the PTEN gene (MOE PTEN^shRNA^), in which case the reduction of PTEN induces tumorigenic phenotype.(24)

MOSE cells were cultured in αMEM (MT10022CV, Fisher) supplemented with 10% fetal bovine serum (FBS, 21G267, Sigma-Aldrich), 2 mM L-glutamine (TCG0063, VWR), 10 mg/mL ITS (11074547001, Sigma-Aldrich), 1.8 ng/mL EGF (100-15, Peprotech Inc.), 100 U/mL penicillin-streptomycin (15140-122, Gibco), and 1 mg/mL gentamycin (30-005-CR, CellGro). The MOE cell lines (SCR^shRNA^ and PTEN^shRNA^) were maintained in similar media but with the addition of 18.2 ng/mL estradiol-17β (E1024, Sigma-Aldrich) and selection antibiotics.

### 3. Coculture incubation for downstream MSI analyses

Low melting agarose (2%, A9414, Sigma-Aldrich) was liquified from a solid at 70□. Cells maintained in αMEM media in T-75 flasks were rinsed with PBS (16777, VWR) and detached using 1 x trypsin (25200072, Life Technologies). Detached cells were transferred to 10 mL αMEM media and counted using an automated cell counter (Accuris E7500 QuadCount™). Aliquots of cells in αMEM were transferred to centrifuge tubes and centrifuged for 5 minutes at 900 rpm (Eppendorf 5804 R Benchtop Centrifuge). After removing the αMEM media, cell pellets were resuspended in 2x DMEM media (D5523, Sigma-Aldrich) supplemented with 10% FBS and 2x penicillin-streptomycin to a concentration of 333 cells/µL. These cell suspensions were mixed 1:1 with 2% agarose to yield a final cell concentration of 166 cells/µL in 1% agarose and 1x DMEM media.

Halved omental explants were placed in the corner of wells in an 8-well chamber (177445, Lab-Tek) attached to an ITO-coated glass slide (8237001, Bruker Daltonics). Each condition (300 µL) was plated into a well, ensuring the tissue remained in the corner of the well. For the divided chamber layout, a removable plastic divider was placed diagonally across each well.(20,21) Cells in agarose were plated first (150 µL) on one side of the divider. After the agarose solidified, the divider was removed, and tissue in agarose (150 µL) was plated on the other side.

Slides were placed in a humidified incubator at 37°C and 5% CO_2_ and cocultured for 4 days. After 4 days, the 8-well chamber was detached from the glass slide, and omental tissue was removed with a razor blade prior to desiccation. Agarose cocultures were desiccated on the ITO-glass slide in an incubator at 30□ on a home-built spinning apparatus for 4 hours.(25)

### 4. Matrix application

MALDI matrices α-cyano-4-hydroxycinnamic acid (CHCA (98%), C2020, Sigma-Aldrich) and 2,5-dihydroxybenzoic acid (DHB (98%), 149357, Sigma-Aldrich) were recrystallized in-house as previously described.(21) The MALDI matrix used for MSI was a 50:50 mixture of CHCA:DHB at 10 mg/mL dissolved in 90:10 ACN:H_2_O with 0.1% TFA applied using a TM sprayer™ (HTX Imaging).

### 5. MSI analysis

Before MSI analysis, slides were scanned using Tissue Scout (Bruker Daltonics) for the initial screen and Epson Perfection V850 Pro for subsequent experiments. Scanned images were used to guide irradiation. MSI data was acquired using timsControl v2.0.51.0_9669_1571 and flexImaging 5.1 software for the initial screen, fleximaging 7.4 for subsequent experiments at 100 μm spatial resolution on a timsTOF fleX mass spectrometer (Bruker Daltonics). Data were collected using a mass range of 50-1500 Da in positive ion mode with the laser width set to 100 μm imaging and the laser power set to 90%. At each raster point, 1,000 laser shots were delivered at a frequency of 1,000 Hz. The instrument was calibrated manually using phosphorus red prior to imaging. MSI data was analyzed, and statistical analysis was performed using SCiLS™ Lab version 2023c core (Bruker Daltonics). All spectra were normalized to the root mean square (RMS). These data are available at MassIVE under accession number MSV000095459.

### 6. Extraction of 3D cocultures for signal validation and quantification

Cocultured agarose plugs were dried *in vacuo* and macerated with a sterile toothpick prior to extraction with 4 mL 50:50 DMF:H_2_O with 0.1% FA. Samples were sonicated for 1 hour before the extract was filtered through a 0.2 µM nylon filter (09719C, Fisher), and the supernatant was dried *in vacuo*.

### 7. MALDI-MS/MS of coculture extracts

Coculture extracts were normalized by dry weight and resuspended in 50:50 MeOH:H_2_O at 1 mg/mL. Valine analytical standard (BP397, Fisher Biotech) was prepared at a concentration of 0.1 mg/mL in 50:50 MeOH:H_2_O. Extracts and analytical standards were mixed 1:1 with CHCA matrix (40 mg/mL in 78:22 ACN:H_2_O with 0.1% TFA) and spotted on an MTP target plate (8280784, Bruker Daltonics). MALDI-MS/MS data was acquired using timsControl v2.0.51.0_9669_1571 on a timsTOF fleX mass spectrometer (Bruker Daltonics). Data were collected using a mass range of 50-650 Da in positive ion mode with the laser width set to 100 µM using an M5 defocus laser. The laser power was set to 60%. At each raster point, 200 laser shots were delivered at a frequency of 1,000 Hz. The collision energy (CE) used for fragmentation was 20 eV. The instrument was calibrated manually using phosphorus red prior to analysis. Data were analyzed using DataAnalysis version 6.1 (Bruker Daltonics).

### 8. LC-MS analysis using aTRAQ kit to quantify amino acids

Coculture extracts were resuspended in H_2_O at 1 mg/mL, and amino acids were quantified using an aTRAQ kit for amino acid analysis of physiological fluids (Sciex). Coculture extracts (10 µL) were derivatized using the protocol and reagents specified by the kit, dried *in vacuo*, and resuspended in 30 µL H_2_O.(26,27) Reverse-phase LC was performed on an Elute UPLC (Bruker Daltonics) using an amino acid analyzer C18 column (5 µm, 4.6 mm × 150 mm, 4374841, Sciex) with a sample injection volume of 2 µL. The mobile phase consisted of A (H_2_O with 0.1% FA and 0.01% hepatofluorobutyric acid (HFBA)) and B (ACN with 0.1% FA and 0.01% HFBA) with a flow rate of 0.4 mL/min. The gradient began with 2% B and was linearly increased to 40% B over 12 minutes; 40% B was held for 8 minutes, then 40% B was linearly increased to 90% B over 2 minutes. The column was washed with 90% B for 2 minutes before linearly decreasing to 2% B over 2 minutes. The column was re-equilibrated with 2% B for 10 minutes. The temperature of the column oven was 50□. MS spectra were collected using a timsTOF fleX mass spectrometer (Bruker Daltonics) in positive ion mode with a mass range of 50-850 Da and a spectra rate of 4 Hz. Prior to analysis, the instrument was calibrated using 0.5 mM sodium formate. All samples were analyzed in three biological replicates (N=3). A normalized response was calculated using the ratio of the area under the curve (AUC) of the analyte peak to the internal standard peak. The concentration of amino acids in coculture extracts was determined by multiplying this normalized response by the concentration of the corresponding internal standard for each amino acid. Since samples were diluted 3x prior to analysis, this result was multiplied by 3 to determine concentrations in the original 1 mg/mL extract. These data are available at MassIVE under accession number MSV000095455.

### 9. Proliferation assay

Cells were plated at a density of 1000 cells per 100 µL in 96-well plates. Cells were allowed to attach to the plate for 24 hours prior to treatment. Plates were fixed with 20% Trichloroacetic acid on days 0, 1, 3, and 5. Cell viability was then determined using 0.04% sulforhodamine B (SRB) via colorimetric detection at 505 nm.(28) Data was normalized to day 0.

### 10. Immunoblotting

250,000 cells were seeded into 6 well plates and left to grow for 24 hours prior to treatment. Cells were removed via trypsin-EDTA, spun down, and resuspended for lysis in RIPA buffer (50 mM Tris pH 7.6, 150mM NaCl, 1% Triton X-100, 0.1% SDS) with protease (04693159001, Roche) and phosphatase inhibitors (524625, Millipore Sigma). Protein concentration was determined by Bradford (5000006, Bio-Rad). Protein lysate (30 μg) was loaded onto an SDS-PAGE gel and transferred to a nitrocellulose membrane. Blots were blocked with 5% BSA in TBS-T and probed at 4°C overnight with primary antibodies, washed thrice, incubated with secondary antibody for 30 minutes, washed thrice, and then developed as described previously.(29) Primary antibodies were utilized at a concentration of 1:1000 and included the following: mTOR (CST 2983), p-mTOR (CST 2971), p70S6K (CST 9202), p-p70S6K (CST 9234) and GAPDH (CST 2118). Secondary antibody was anti-rabbit and HRP-linked (7074S, Cell Signaling) and used at a concentration of 1:10,000.

## RESULTS

### MSI coculture adaptation for use with omental tissue

While our MSI method was originally envisioned as a plug-and-play technique that could be adapted to many biological cell cultures and organoid cultures, adaptation of this protocol for use with omental tissue was necessary. Matrix-assisted laser desorption ionization (MALDI)- MSI requires a flat sample (30,31), and due to its fatty nature, omental tissue does not dry flat but crystalizes in heat when desiccated (Figure S1). To address this inconsistent omental crystallization, we placed the tissue in the corner of the 8-well chamber, rather than the center as we have previously reported with murine ovary explants, and removed the omental tissue from cocultured agarose plugs prior to sample desiccation and MSI analysis (Figure 1A). We confirmed that the omental tissue is viable after four days of incubation in agarose (Figure S2).

**Figure 1:**
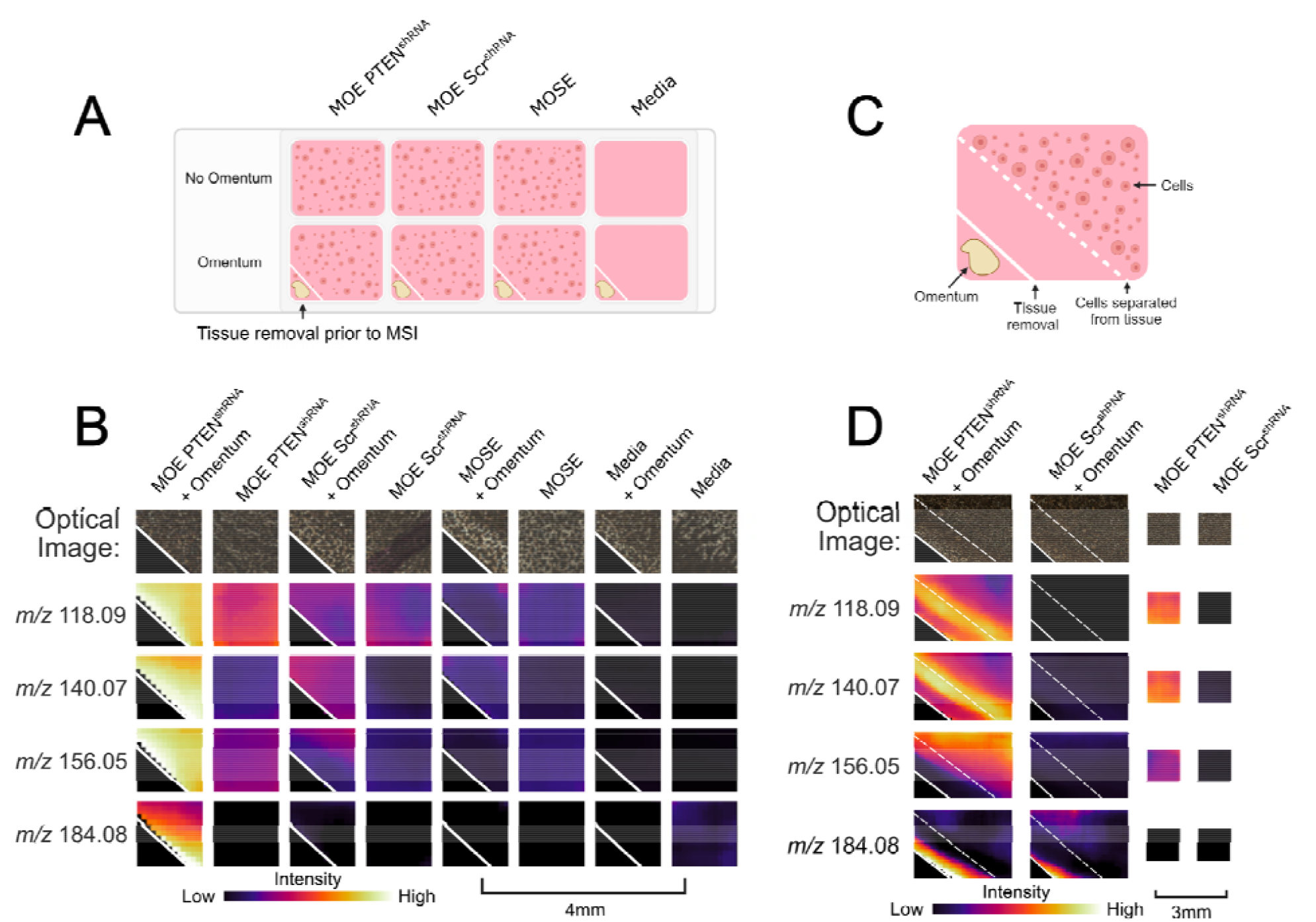
MSI analysis shows several signals specific to tumorigenic FTE cell/omentum coculture condition. **A)** Illustration of imaging layout on the slide for untargeted analysis which includes tumorigenic FTE cells (MOE PTEN^shRNA^) and non-tumorigenic controls. **B)** Ion images of 4 representative signals upregulated in the HGSOC condition specific to the PTEN mutation (N=3). **C)** Illustration of divided well layout to identify the origin of signals. **D)** Signals in B are replicated when cells and omentum are physically separated in agarose. Signals at *m/z* 118, *m/z* 140, and *m/z* 156 originate from tumorigenic FTE cells, while the signal at *m/z* 184 originates from omental tissue. These signals are statistically significant compared to MOE PTEN^shRNA^ + omentum (p < 0.05).

### Initial MSI screen revealed several signals increased in tumorigenic FTE cell/omentum coculture, and the origin of these signals

We chose to utilize a murine oviductal epithelial (MOE) cell line with an shRNA targeting the PTEN gene (MOE PTEN^shRNA^) as our tumorigenic FTE cell model since we have previously demonstrated that silencing PTEN was sufficient to drive colonization of both the ovary and the omentum in vivo using athymic mice.(32) This model is advantageous because the omental tissue and FTE cells are from the same species, eliminating species differences. In the experimental design for the initial screen, we included several controls, including a MOE cell line expressing a scrambled control shRNA (MOE SCR^shRNA^) to test for cells that are not tumorigenic, a murine ovarian surface epithelial (MOSE) cell line to test for cell specificity, and a media negative control. Each condition was cultured alone and cocultured with omental tissue (Figure 1A). Samples were incubated for 4 days to allow interaction to occur prior to imaging (N=3). Samples were analyzed using a MALDI-quadrupole time-of-flight (MALDI-QqTOF) mass spectrometer.

The original MSI screen yielded over 25 signals specific to the tumorigenic FTE cell/omentum coculture condition, of which four representative signals are depicted in Figure 1B and replicated in Figure S3-S4. These signals were determined to be significantly increased (p <0.05) using the MSI data analysis software SciLS when comparing MOE PTEN^shRNA^ + omentum against all other conditions including scrambled + omentum.(33) Additional MSI using a divided chamber coculture format (Figure 1C) revealed the origin of signals and confirms signals that accumulate due to secretion in agarose without direct physical contact between the cells and the tissue. Three signals (*m/z* 118, *m/z* 140, and *m/z* 156) were found to be produced by tumorigenic FTE cells alone, while one signals (*m/z* 184) wase found to be produced by omental tissue based on the spatial distribution (Figure 1D and S5). Signals were putatively annotated by searching for metabolites that matched the experimental monoisotopic mass in the Human Metabolome Database (HMDB, Table 1).(34,35) It is worth noting that all assignments are designated as putative due to the nature of the ppm error achieved from our MSI samples. It is a well-documented effect that the slight surface heterogeneity and added height from the glass slides contributes to the higher mass errors observed. Putative annotations for these signals suggested alterations in amino acid metabolism and the potential induction of catecholamine signaling in cocultures (Table 1). Valine and histidine are both essential amino acids and epinephrine is a catecholamine neurotransmitter.(36)

**Table 1:** Putative annotations and ppm error. Signals (*m/z*) identified in the initial MSI screen and putative annotations based on the adduct and searching the expected monoisotopic mass in the human metabolome database (HMDB).(36) Ppm error was calculated based on the measured accurate mass and the calculated exact mass of putatively annotated metabolites in their observed adduct form.

| MSI Signal (Observed) | Putative Adduct | Exact mass (calc) | Putative annotation | ppm error |
| --- | --- | --- | --- | --- |
| *m/z* 118.0918 | [M+H]<sup>+</sup> | 118.0863 Da | Valine | 46.6 |
| *m/z* 140.0735 | [M+Na]<sup>+</sup> | 140.0682 Da | Valine | 37.8 |
| *m/z* 156.0470 | [M+H]<sup>+</sup> | 156.0768 Da | Histidine | -190.99 |
| *m/z* 184.0775 | [M+H]<sup>+</sup> | 184.0968 Da | Epinephrine | -104.8 |

### Valine annotation was confirmed using tandem mass spectrometry

While we were able to identify spatially interesting signals and determine their relative abundance using MSI, we were unable to obtain high-quality fragmentation patterns directly from the MSI sample due to low overall signal intensity and ion suppression within the agarose culture. To obtain fragmentation patterns, metabolites were extracted from agarose cocultures using organic solvents, and the extracts were resuspended at 1 mg/mL (Figure S6A). Extracts were then analyzed alongside commercial standards of putatively identified metabolites using the dried droplet method for MALDI-MS/MS (Figure S6A). The fragmentation pattern for *m/z* 118 in the tumorigenic FTE cell/omentum coculture matched the fragmentation pattern from L-valine (Figure S6B), thus confirming the putative annotation with level 2 confidence.(37,38)

### Amino acids quantified using LC-MS showed BCAA catabolism is increased in tumorigenic FTE cell/omentum cocultures

To orthogonally validate putative annotations and further probe alterations in amino acid metabolism, amino acids in coculture extracts (N=3) were quantified using an aTRAQ kit (Sciex) with LC-MS analysis.(26,27) Overall, we could detect and quantify 24 amino acids by retention time matching with internal standards. Thus, each amino acid was identified with level 1 confidence (Table S1).(37) Serine was significantly decreased (p < 0.05) in the tumorigenic FTE cell/omentum condition compared to the DMEM control. (Figure 2D). Interestingly, data regarding valine and other BCAAs, including leucine and isoleucine, revealed decreased levels in the tumorigenic FTE cell/omentum condition compared to media alone (Figure 2A-C). (39) The aTRAQ kit is capable of quantifying 44 different amino acids and amino acid derivatives; of these, 24 were detected and quantified in coculture extracts, and 20 were not detected (Table S2 and Figure S7).

**Figure 2:**
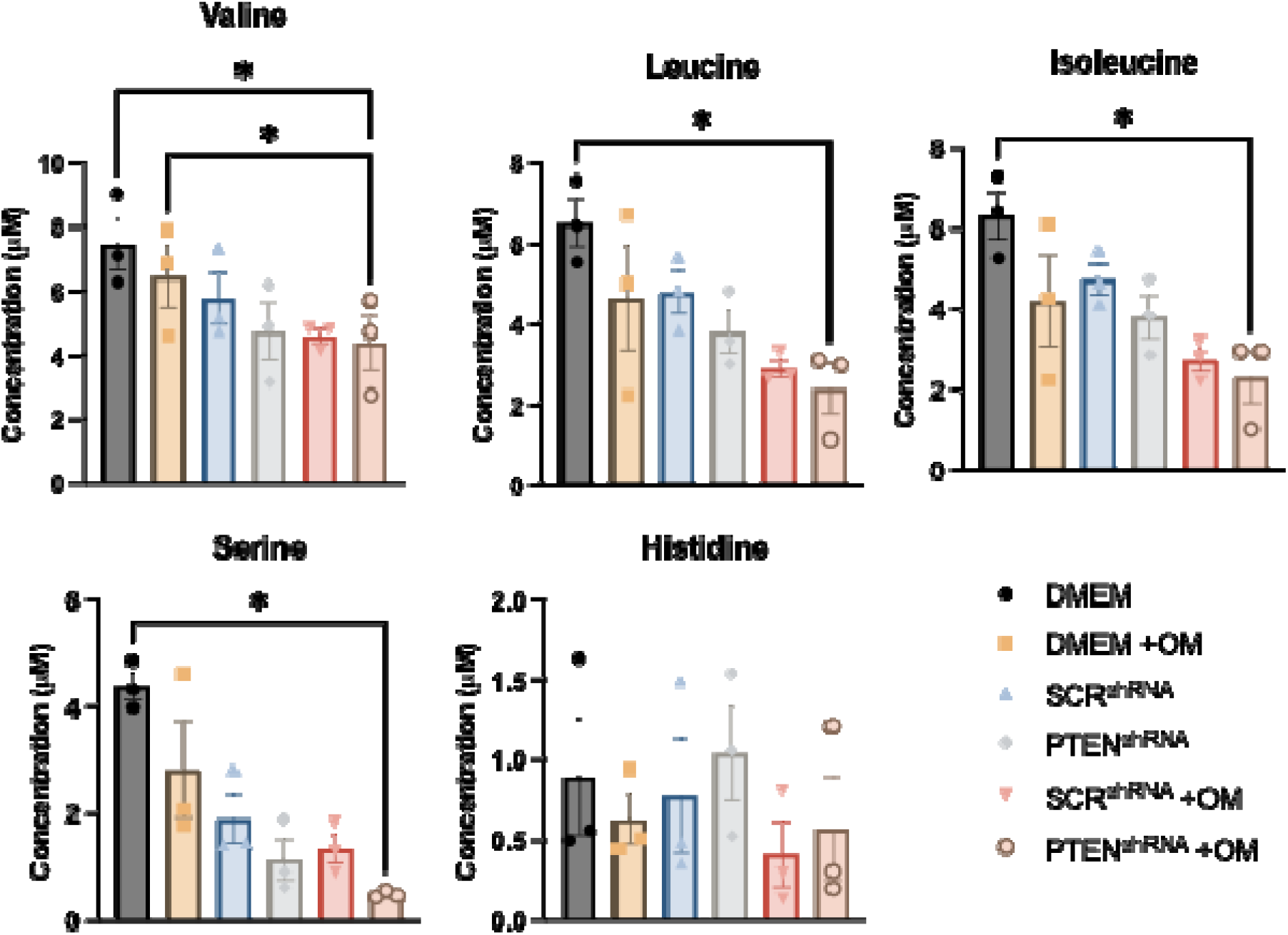
LC-MS data shows increased consumption of BCAA’s and serine, a trend toward increasing glutamic acid and no change in histidine concentration. LC-MS results from the analysis of 3D agarose coculture extracts using aTRAQ kit to determine concentrations of **A)** valine, **B)** isoleucine, **C)** leucine, and **D)** serine **E)** histidine in extracts resuspended at 1 mg/mL. Significance was determined using a one-way ANOVA with Tukey’s post hoc and a cutoff value of 0.05.

### Analysis of the valine calibration curve using MSI shows evidence of ion suppression

Based on the aforementioned results, we tested the hypothesis that ion suppression caused the apparent ‘increase’ in valine levels in tumorigenic FTE/omentum cocultures (Figure 1B, Figure S3), while our quantitative LC-MS data showed a decrease (Figure 2A). Ion suppression is a type of matrix effect where certain metabolites are inhibited from being ionized and, consequently, remain undetected. This phenomenon can arise due to various factors, including the presence of highly abundant metabolites that compete for ionization.(40) As a result, some metabolites may exhibit a linear response, with their signal intensity increasing as concentrations rise until reaching a threshold. Beyond that threshold, the signal intensity begins to decrease with further increases in concentration.

The media used for mammalian cell culture is supplemented with high concentrations of amino acids, including L-valine at a concentration of 800 µM. We hypothesized that this high concentration of L-valine may induce ion suppression, and the consumption of L-valine in tumorigenic FTE/omentum cocultures may alleviate ion suppression, thereby leading to increased signal observed in MSI. We tested this hypothesis using standard curves with known concentrations of a valine analytical standard spiked into agarose. The first standard curve had concentrations of 800 µM, 400 µM, 80 µM, 8 µM, 0.8 µM, and 0.08 µM valine. MSI analysis revealed that no signal was observed from 800 µM to 80 µM, and the highest signal intensity for *m/z* 118 was observed at the 8 µM concentration. Below the 8 µM concentration, the signal intensity dropped off, a result that is consistent with our hypothesis (Figure 3A). For further confirmation, we analyzed a second standard curve with a 4 x serial dilution of 20 µM valine. Again, the signal intensity was low at the 20 µM concentration, peaked at the 5 µM concentration, and decreased at concentrations lower than 5 µM. The intermediate concentrations, at 10 µM and 2.5 µM, showed similar signal intensities for *m/z* 118, providing additional confidence that our hypothesis was confirmed (Figure 3A). We noted that histidine also had a similar aTRAQ trend but was not significantly altered and subsequently does not suffer from ion suppression (Figure 3B).

**Figure 3:**
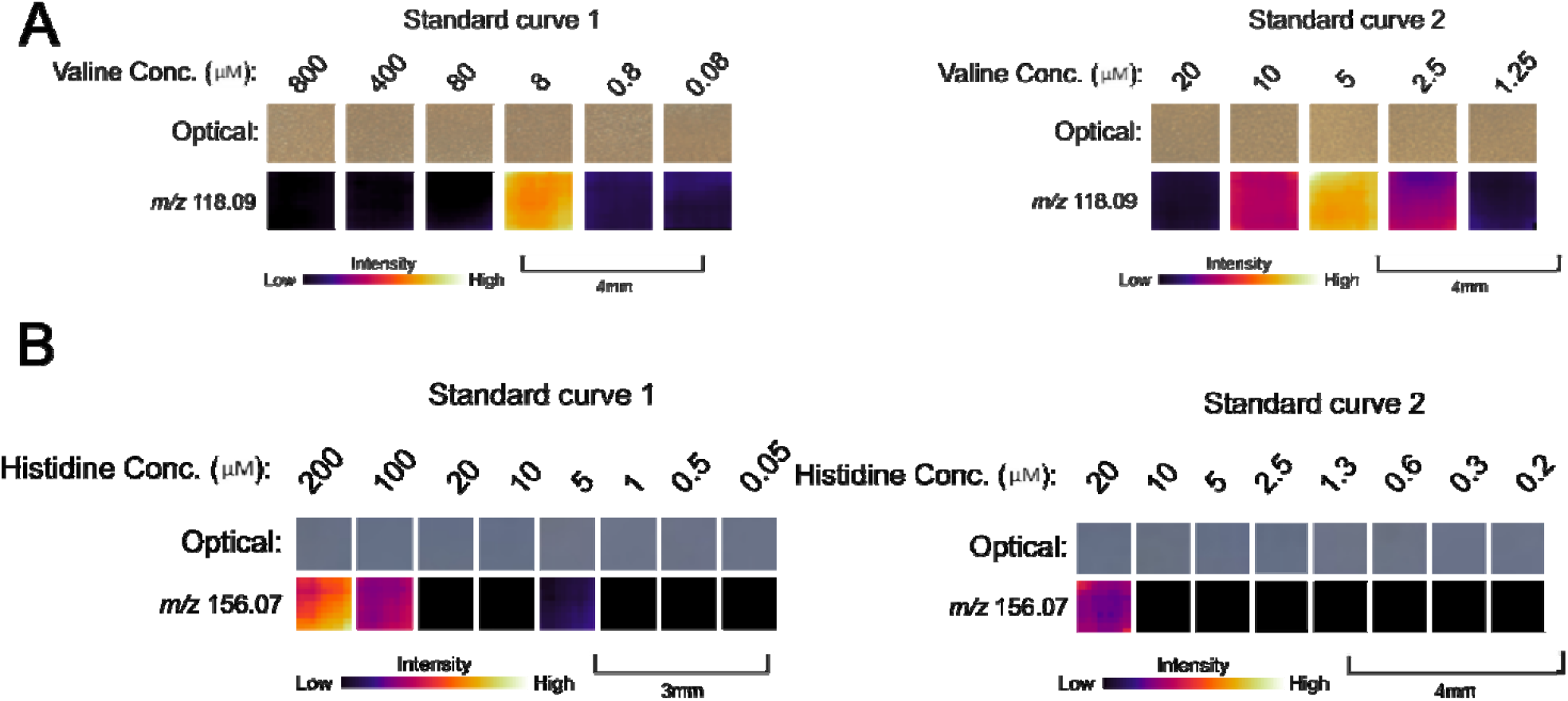
Ion suppression tests for select amino acids in cell culture media. **A)** Ion images showing signal *m/z* 118.09 with different concentrations of a valine analytical standard spiked into agarose. Standard curve 1 shows concentrations ranging from 800 µM to 0.08 µM, while standard curve 2 shows concentrations ranging from 20 µM to 1.25 µM. DMEM media contains 800 µM L-valine. **B)** Ion images showing signal *m/z* 156.07 with different concentrations of a histidine analytical standard spiked into agarose. Standard curve 1 shows concentrations ranging from 200 µM to 0.005 µM, while standard curve 2 shows concentrations ranging from 20 µM to 0.2 µM. DMEM media contains 200 µM L-histidine.

### Valine supplementation leads to increased proliferation and phosphorylation of mTOR in tumorigenic FTE cell cultures

BCAAs are essential amino acids that mammalian cells cannot synthesize, so cancer cells likely acquire them through protein degradation or from the tumor microenvironment (TME). These amino acids play a fundamental role as building blocks for protein synthesis. They can be metabolized into branched-chain □-keto acids (BCKAs) within the cytosol by branched-chain amino acid transaminase 1 (BCAT1) or within the mitochondria by branched-chain amino acid transaminase 2 (BCAT2).(39) This process involves the conversion of α-ketoglutarate (α-KG) to glutamate. BCAAs may also serve as nitrogen sources for the biosynthesis of nucleotides and non-essential amino acids through the glutamate-glutamine axis.(39) Additionally, they can be catabolized to produce acetyl-CoA and succinyl-CoA, which are utilized in the TCA cycle, thereby contributing to energy production.(39) The BCAA leucine is a well-known mechanistic target of rapamycin (mTOR) agonist, and studies indicate that other BCAA’s may also be capable of activating mTOR signaling.(41,42) mTOR functions, in part, by regulating the phosphorylation of p70 S6 kinase (p70S6k).(43) This signaling pathway regulates proliferation and is often activated in tumors.(44) We sought to characterize the impact of valine supplementation on proliferation and mTOR signaling in tumorigenic FTE cells, as BCAAs are known to stimulate mTOR and proliferation in other cell types (Figure 4A). We evaluated the proliferation in media supplemented with 800 µM (0.8 mM, standard DMEM concentration) and 1.6 mM valine. We evaluated the phosphorylation of mTOR in 1.6 mM valine-supplemented media after 24 hours by western blot. The results show that the phosphorylation of mTOR increased with 1.6 mM valine supplementation (Figure 4B-C), indicating that valine can stimulate the phosphorylation of mTOR in tumorigenic FTE cells. We also found that there was a significant increase in proliferation of tumorigenic MOE PTEN^shRNA^ cells (at day 5) and MOE PTEN^shRNA^ p53^R273H^ cells (day 3) when treated with 1.6 mM valine-supplemented media compared to DMEM (Figure 4D). This data indicates that 1.6 mM valine supplementation increases the proliferation of tumorigenic FTE cells.

**Figure 4:**
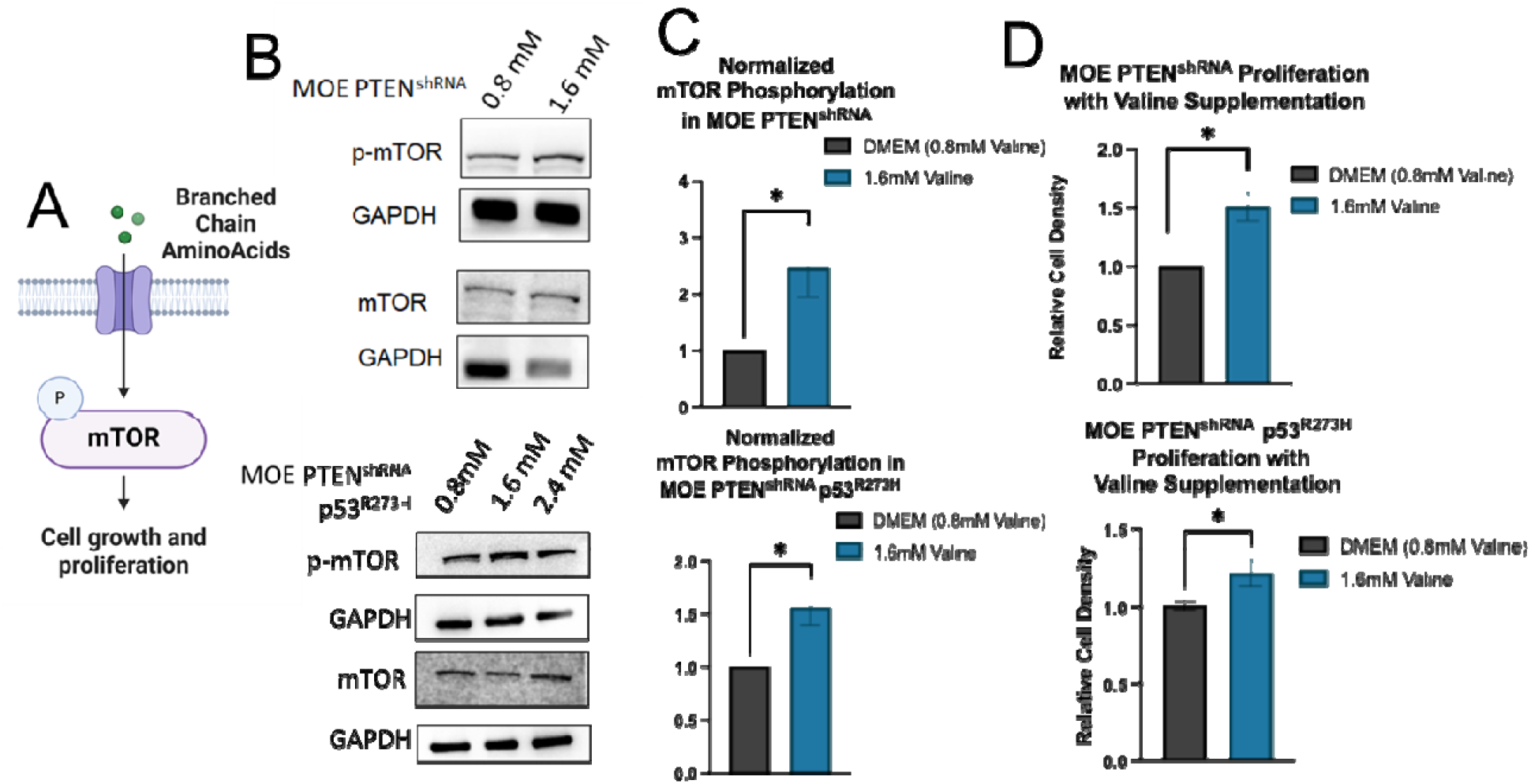
Valine supplementation increases the proliferation of tumorigenic FTE cells and activates mTOR signaling. **A)** Schematic of mTOR activation by BCAA. **B)** Representative western blot showing increased phosphorylation of mTOR with 1.6 mM valine supplementation in MOE PTEN^shRNA^ and MOE PTEN^shRNA^ p53^R273H^. **C)** Quantification of western blot (N=3). **D)** SRB proliferation data from MOE PTEN^shRNA^ and MOE PTEN^shRNA^ p53^R273H^ cells supplemented with 1.6 mM valine supplementation.

## DISCUSSION

In this study, we adapted our MSI protocol to investigate metabolic alterations at the interface of tumorigenic FTE cells and omental tissue to elucidate the chemical cues involved in secondary metastasis. Our analysis showed that several signals were elevated in tumorigenic FTE cell/omentum cocultures compared to the controls. Based on our initial screens, putative annotations suggested that amino acid metabolism was perturbed at this coculture interface. Notably, the signal at *m/z* 118 was confirmed to represent the BCAA valine, and subsequent quantitative LC-MS analysis revealed increased BCAA catabolism. We attributed the discrepancy between MSI and LC-MS results to ion suppression resulting from high concentrations of valine in the media. Furthermore, proliferation assays demonstrated that valine supplementation promotes the proliferation of tumorigenic FTE cells, and immunoblotting showed an increase in the phosphorylation of mTOR in two cell lines.

Previous research has highlighted the role of amino acids in interactions between OvC cells and omental tissue.(11,15) In our quantitative assay, both glutamine and asparagine were elevated in conditions involving omental tissue (Figure S7). Since the omentum was removed prior to extraction, the increased presence compared to media alone implies their secretion by omental tissue. Glutamine is a vital nutrient for cancer cell survival, especially under metabolic stress conditions;(45) it has been shown to bolster proliferation by activating mTOR and contribute to resistance to chemotherapy in OvC.(41,46–50) Studies indicate that asparagine can sustain cancer cell survival when glutamine is scarce.(51) Previous research suggests that cancer cells may acquire asparagine from the TME through release by CAFs, and similar to glutamine, asparagine has been shown to stimulate mTOR signaling.(48,51,52) Another intriguing finding is the significant decrease in serine and methionine levels in tumorigenic FTE cocultures compared to healthy FTE controls (Figure S7). Both serine and methionine play roles in one-carbon metabolism, providing 1-carbon units for DNA-histone methylation.(53) Serine is also involved in glutathione biosynthesis, a metabolite previously reported to be increased in the OvC TME.(53–55) Surprisingly, although histidine was identified as a putative MSI signal, it was found to be decreased in omental conditions with no difference in tumorigenic FTE compared to healthy FTE (Figure S7). We did not observe increases in the levels of arginine or citrulline, which a previous study reported being secreted into the OvC TME.(11,15) The most notable discovery from our quantitative LC-MS data is the heightened consumption of valine and other BCAAs, which contradicted the MSI results.

The discrepancy between MALDI-MSI and LC-MS outcomes was ascribed to a phenomenon known as ion suppression. Ion suppression is a well-known phenomenon that can reduce signal in both MALDI-MSI and LC-MS.(40,56) In mass spectrometry, the molecular complexity of a sample affects the desorption and ionization of metabolites within it.(56) At elevated concentrations, certain metabolites hinder their own ionization, causing signal intensity to rise until reaching a threshold concentration, beyond which it decreases. Through valine standard curves, we determined that the increase in signal observed in MSI was due to the relief of ion suppression resulting from valine consumption in tumorigenic FTE/omentum cocultures. By extracting metabolites from omental cocultures and diluting them to 1 mg/mL, we reduced the sample complexity and valine concentration in LC-MS samples, thereby mitigating ion suppression. This discovery holds significant implications for future MSI investigations; when a metabolite of interest is known to have a high concentration in the media, researchers should consider analyzing a standard curve to assess the impact of ion suppression. Furthermore, this underscores the importance of orthogonally validating MSI results.

The catabolism of valine and other BCAAs was observed to be higher in tumorigenic FTE cell/omentum cocultures compared to media alone conditions. Notably, while the difference in BCAA levels between tumorigenic FTE and healthy FTE cocultures was not statistically significant in extracts by LC-MS, MSI data in a divided chamber format revealed a more pronounced effect in cells closely situated to omental tissue. This observation suggests that the increased BCAA catabolism results from an interaction. Additionally, this finding may elucidate why the effect was not significant in coculture extracts of whole agarose plugs, as many cells within the coculture are distant from the omentum and may not experience this interaction. Elevated BCAA catabolism in OvC is substantiated by previous studies demonstrating increased expression of the BCAA catabolic enzyme BCAT1 in HGSOC.(57,58) Knockdown of BCAT1 has been shown to repress growth, supporting its significance in cancer progression.(57,58) In a study by Zhang and Han, elevated levels of BCAT1 were found to stimulate proliferation and mTOR activity in breast cancer.(59) These findings are consistent with our proliferation data, which showed that valine supplementation increased proliferation and the phosphorylation of mTOR in tumorigenic FTE cell cultures. When hyperactivated, mTOR signaling promotes cell proliferation and metabolism that aids in tumor progression; mTOR signaling is enhanced in various types of cancer, and inhibition of this pathway is a promising therapeutic target.(60)

## CONCLUSION

Our study sheds light on the metabolic dynamics implicated in the secondary metastasis of HGSOC to the omentum. Using MSI analysis, we uncovered alterations in amino acid metabolism and observed an increase in BCAA catabolism at the interface of OvC cells and omental tissue. This heightened BCAA catabolism may facilitate increased proliferation of OvC cells through mTOR phosphorylation at the omentum/cancer cell interface. Moving forward, our research will focus on investigating other MSI signals and metabolites identified in this study, as well as delving into the molecular mechanisms underlying the observed increase in BCAA catabolism at this interface. By elucidating the intricate interplay of chemical cues involved in secondary metastasis, our findings enhance our understanding of OvC pathogenesis and hold promise for the development of novel therapeutic strategies targeting the metastatic TME in HGSOC.

## ASSOCIATED CONTENT

### Supporting Information

The supporting information is available free of charge.

## Author Contributions

The manuscript was written with contributions from all authors. All authors have given approval to the final version of the manuscript.

## Acknowledgments

Figure 1A, Figure 1C, and Figure 4A were created with BioRender.com

## Funding Sources

This work is supported in part by the National Institute of Health grant R01 CA240423 (JB and LS), the Laura Crandall Brown Foundation Ovarian Cancer Early Detection Research Grant from the Foundation for Women’s Cancer supported by the Laura Crandall Brown Foundation (to LS and J B). HL was supported by NIH R01 CA240423-03S1, and NIH F30 CA260791 supported TB.

## Abbreviations

OvC: ovarian cancer
HGSOC: high-grade serous ovarian cancer
FTE: fallopian tube epithelial
CAF: cancer-associated fibroblast
TME: tumor microenvironment
MSI: mass spectrometry imaging
BCAA: branched-chain amino acid
AUC: area under the curve
CE: collision energy
MALDI: matrix-assisted laser desorption ionization
MOSE: murine ovarian surface epithelial
MOE: murine oviductal epithelial
Q-TOF: quadrupole time-of-flight
LC-MS: liquid chromatography-mass spectrometry
mTOR: mechanistic target of rapamycin

## Notes

https://massive.ucsd.edu/ProteoSAFe/dataset.jsp?task=da354312aae04cc6a920daea4581341a

https://massive.ucsd.edu/ProteoSAFe/dataset.jsp?task=b824a783b9424099b39e962eb13cf3ca

### Competing Interest Statement

The authors have declared no competing interest.

