## Supplemental Information for "Branched-chain amino acid catabolism promotes ovarian cancer cell proliferation via phosphorylation of mTOR"

**Figure S1.** Omentum coculture drying photo…………………………...………………………….. S2
**Figure S2.** Live dead staining of omental tissue…………………………………………………… S3
**Figure S3.** Omentum IMS replicates ……………………………………………………………..… S4
**Figure S4.** Other signals identified in the initial MSI screen (N=3)…………….…………..…………….. S5
**Figure S5.** Split chamber MSI signals………………………………………………………………. S6
**Table S1.** Putative annotations of other signals identified in the initial MSI screen……………... S7
**Figure S6.** Extraction protocol and MALDI-MS/MS data…………………………………………….. S8
**Table S2.** Ppm errors for amino acids quantified using aTRAQ kit…..…………..……………….. S9
**Figure S7.** LC-MS data from amino acids quantified using aTRAQ kit.…………………………… S10

**Figure S1.** Omentum coculture drying photo. Omental tissue did not dry flat but crystallized when desiccated using heat.

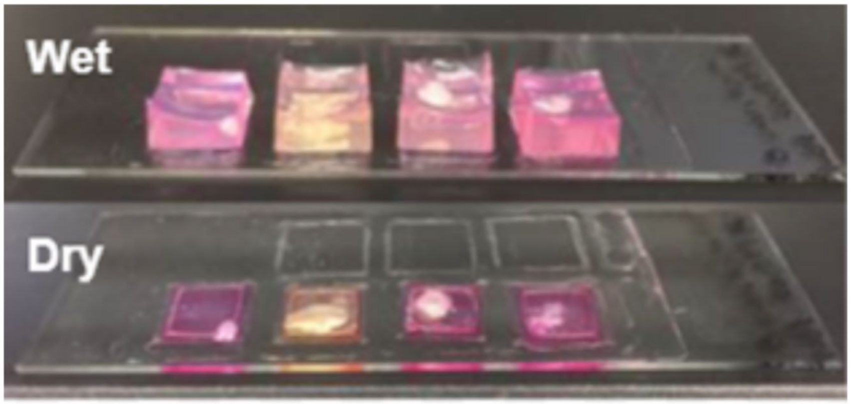

**Figure S2.** Live/Dead staining of omental tissue after 24 hours at 4 degrees and after 4 days at 37 degrees with Calcein AM (Live Green) and BOBO-3 Iodide (Dead Red) .

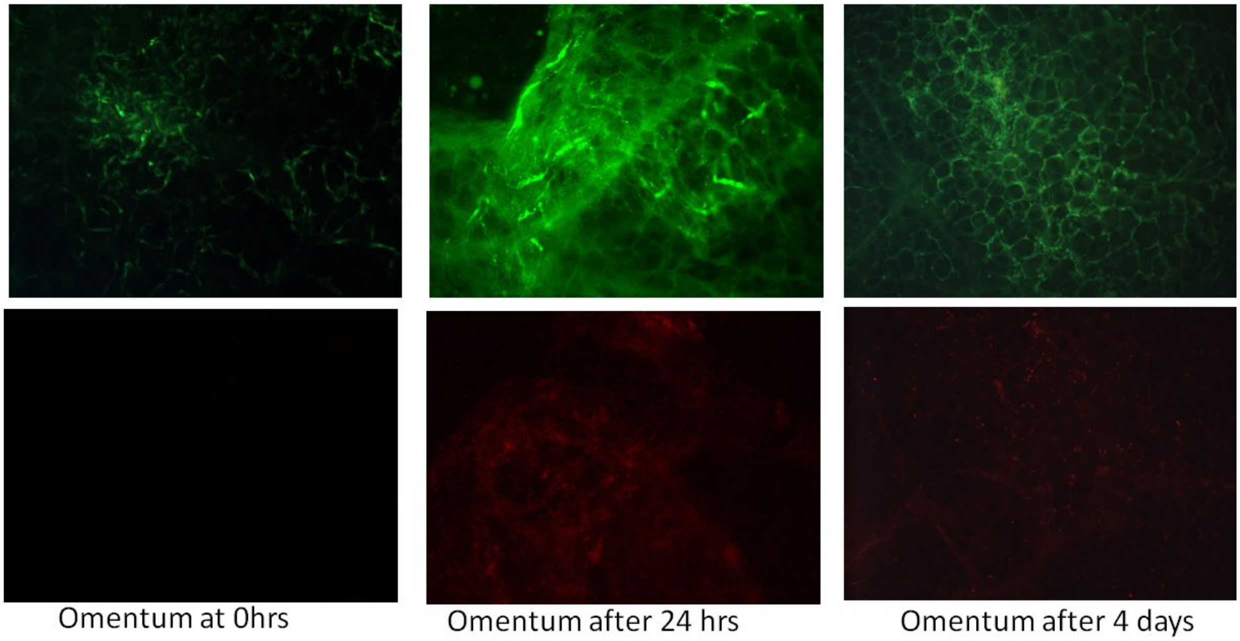

**Figure S3.** Omentum IMS replicates (N=3).

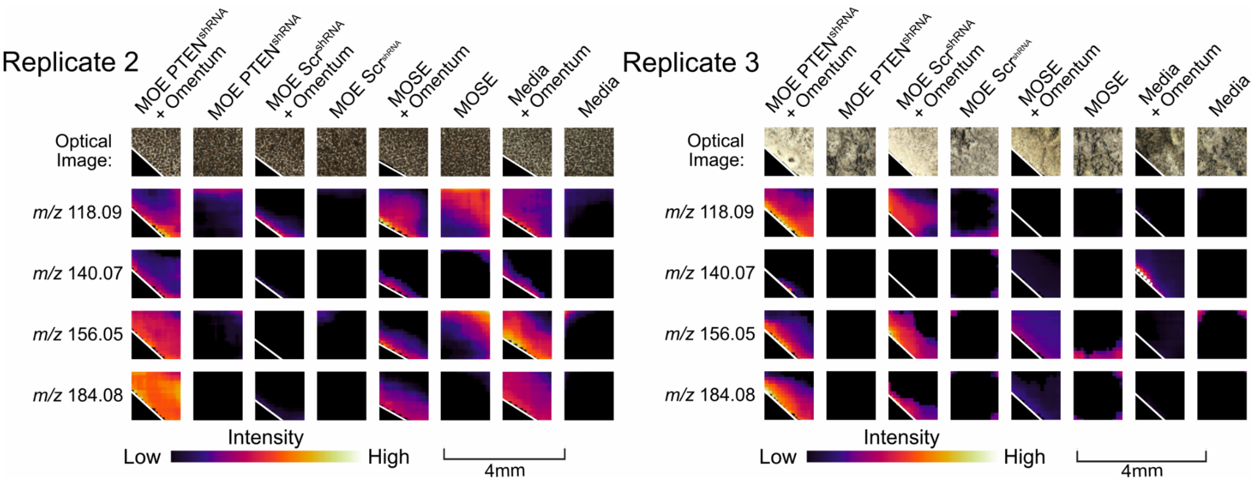

**Figure S4.** Other signals identified in the initial MSI screen (N=3).

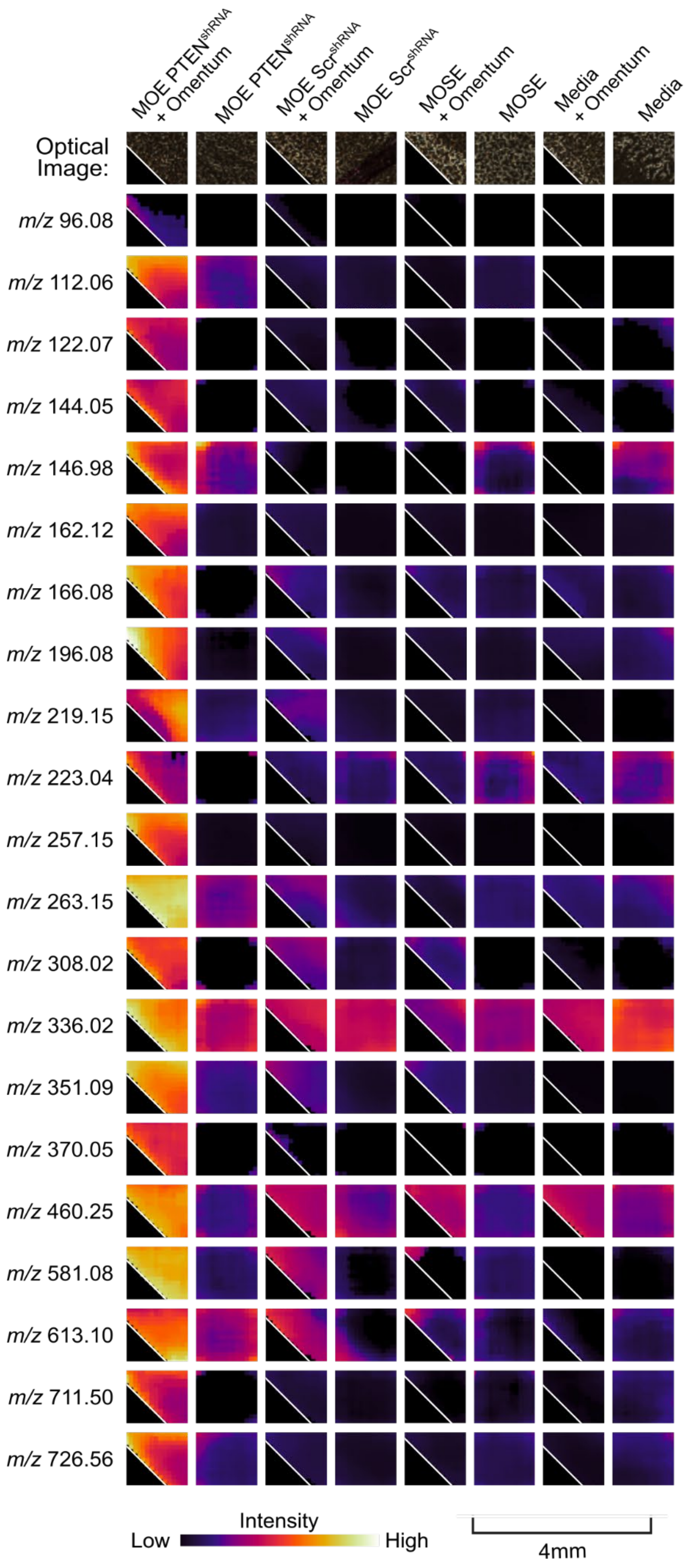

**Figure S5.** Signals from Figure S4 that replicated when cells and omentum are physically separated in agarose. Several signals originate from tumorigenic FTE cells, and several signals originate from omental tissue.

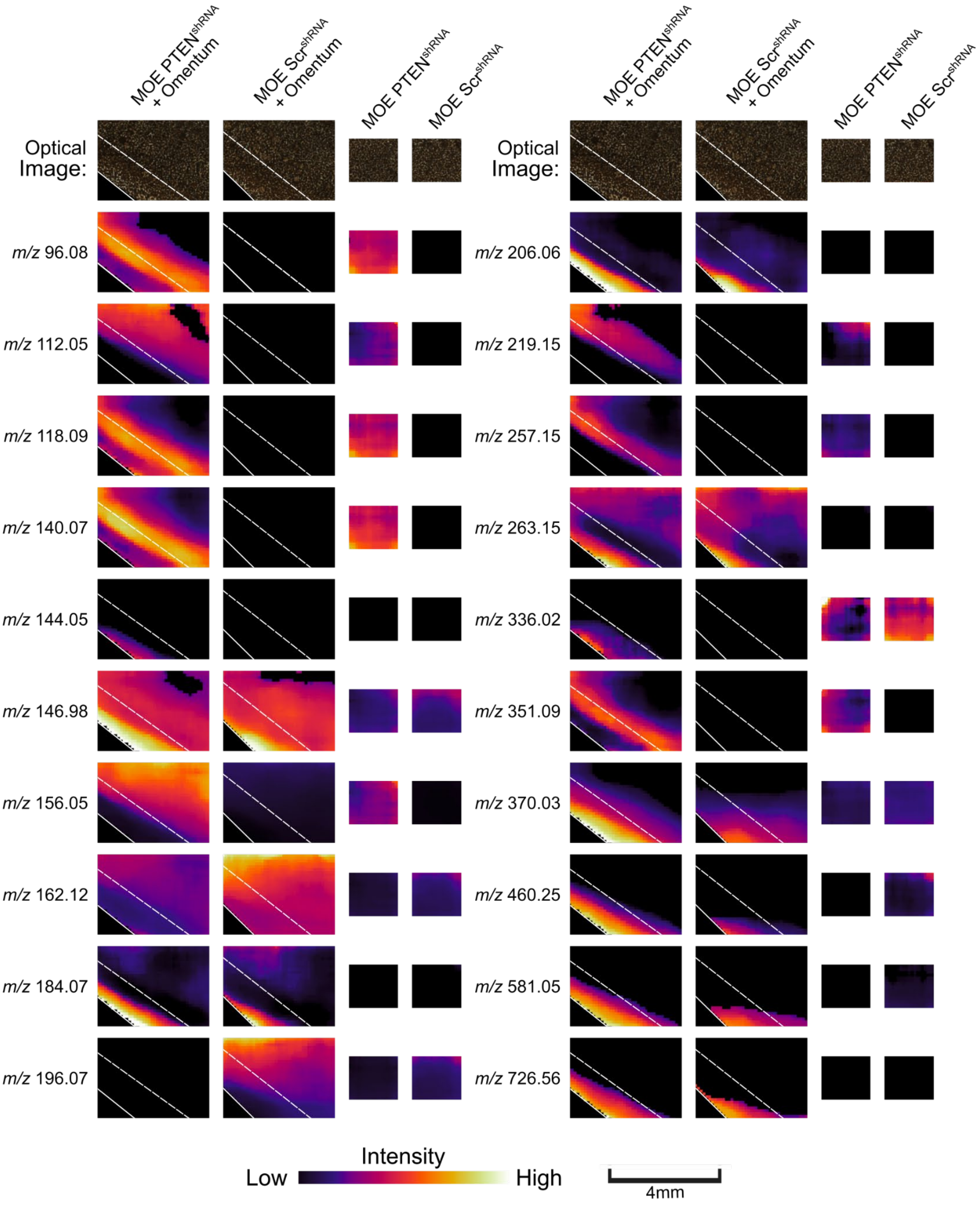

**Table S1.** Putative annotations of other signals identified in the initial MSI screen.

| Signal | Adduct | Putative annotation |
|---|---|---|
| *m/z* 96.08 | $[M+Na]^+$ | 2-Methyl-1-propylamine |
| *m/z* 112.06 | $[M+H]^+$ | Histamine |
| *m/z* 122.07 | $[M+H]^+$ | L-Cysteine |
| *m/z* 144.05 | $[M+H]^+$ | Vinylacetylglycine |
| *m/z* 146.98 | $[M+Na]^+$ | Phosphonoacetaldehyde |
| *m/z* 162.12 | $[M+H]^+$ | L -Carnitine |
| *m/z* 166.08 | $[M+H-H_2O]^+$ | Epinephrine |
| *m/z* 196.08 | $[M+H]^+$ | Dopaquinone |
| *m/z* 219.15 | $[M+H]^+$ | N-acetylserotonin |
| *m/z* 223.04 | $[M+H]^+$ | Cystathionine |
| *m/z* 257.15 | $[M+K]^+$ | N-acetylserotonin |
| *m/z* 263.15 | $[M+H]^+$ | Methylmalonylcarnitine |
| *m/z* 308.02 | $[M+H]^+$ | Deoxycytidine monophosphate (dCMP) |
| *m/z* 336.02 | $[M+H]^+$ | Dihydroneopterin phosphate |
| *m/z* 351.09 | $[M+H]^+$ | Estrone sulfate |
| *m/z* 370.05 | $[M+Na]^+$ | Adenosine monophosphate (AMP) |
| *m/z* 460.25 | $[M+H]^+$ | N-Docosahexaenoyl Methionine |
| *m/z* 581.08 | $[M+H]^+$ | Uridine diphosphate glucuronic acid |
| *m/z* 613.10 | $[M+H]^+$ | Oxidized glutathione |
| *m/z* 711.50 | $[M+H]^+$ | Phosphatidic acid (PA) |
| *m/z* 726.56 | $[M+H]^+$ | Phosphatidylethanolamine (PE) or Phosphatidylcholine (PC) |

**Figure S6.** MALDI-MS/MS fragmentation data from tumorigenic FTE/omentum coculture extract reveals the signal at *m/z* 118 represents L-valine. **A)** Illustration of extraction procedure and MALDI-MS/MS analysis of tumorigenic FTE/omentum coculture extract **B)** Butterfly plot showing fragmentation patterns obtained from MALDI-MS/MS analysis of *m/z* 118 in the tumorigenic FTE/omentum coculture extract (top) and an L-valine analytical standard (bottom). The matching fragmentation pattern at 20 eV indicates that the IMS signal at *m/z* 118 represents L-valine.

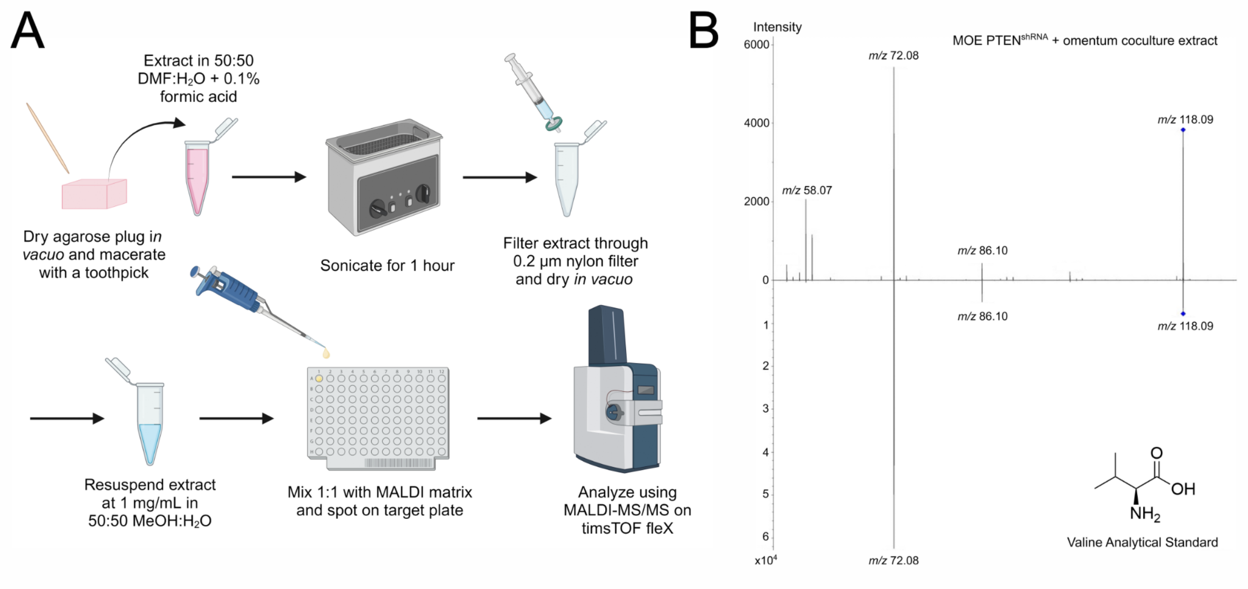

**Table S2.** Ppm errors for amino acids quantified using aTRAQ kit

| Amino acid | Exact mass (calc) | Measured accurate mass | Error |
|---|---|---|---|
| L-alanine | *m/z* 238.1641 | *m/z* 238.1620 | 8.8 ppm |
| L-arginine | *m/z* 323.2281 | *m/z* 323.2255 | 8.0 ppm |
| L-asparagine | *m/z* 281.1699 | *m/z* 281.1679 | 7.1 ppm |
| L-aspartic acid | *m/z* 282.1539 | *m/z* 282.1514 | 8.9 ppm |
| β-Alanine | *m/z* 238.1641 | *m/z* 238.1619 | 9.2 ppm |
| L-citrulline | *m/z* 324.2121 | *m/z* 324.2096 | 7.7 ppm |
| L-cystine | *m/z* 537.2495 | *m/z* 537.2441 | 10.1 ppm |
| Ethanolamine | *m/z* 210.1692 | *m/z* 210.1678 | 6.7 ppm |
| L-glutamic acid | *m/z* 296.1696 | *m/z* 296.1671 | 8.4 ppm |
| L-glutamine | *m/z* 295.1856 | *m/z* 295.1834 | 7.5 ppm |
| Glycine | *m/z* 224.1485 | *m/z* 224.1465 | 8.9 ppm |
| L-histidine | *m/z* 304.1859 | *m/z* 304.1833 | 8.5 ppm |
| Hydroxy- L-proline | *m/z* 280.1747 | *m/z* 280.1724 | 8.2 ppm |
| L-isoleucine | *m/z* 280.2111 | *m/z* 280.2090 | 7.5 ppm |
| L-leucine | *m/z* 280.2111 | *m/z* 280.2089 | 7.9 ppm |
| L-lysine | *m/z* 443.3311 | *m/z* 443.3273 | 8.6 ppm |
| L-methionine | *m/z* 298.1675 | *m/z* 298.1651 | 8.0 ppm |
| L-phenylalanine | *m/z* 314.1954 | *m/z* 314.1929 | 8.0 ppm |
| L-proline | *m/z* 264.1798 | *m/z* 264.1773 | 9.5 ppm |
| L-serine | *m/z* 254.1590 | *m/z* 254.1569 | 8.3 ppm |
| L-threonine | *m/z* 268.1747 | *m/z* 268.1726 | 7.8 ppm |
| L-tryptophan | *m/z* 353.2063 | *m/z* 353.2027 | 10.2 ppm |
| L-tyrosine | *m/z* 330.1903 | *m/z* 330.1875 | 8.5 ppm |
| L-valine | *m/z* 266.1954 | *m/z* 266.1933 | 7.9 ppm |

**Figure S7.** LC-MS data from amino acids quantified using aTRAQ kit (Sciex). Significance was determined using a one-way ANOVA with Tukey's post hoc and a cutoff value of 0.05.

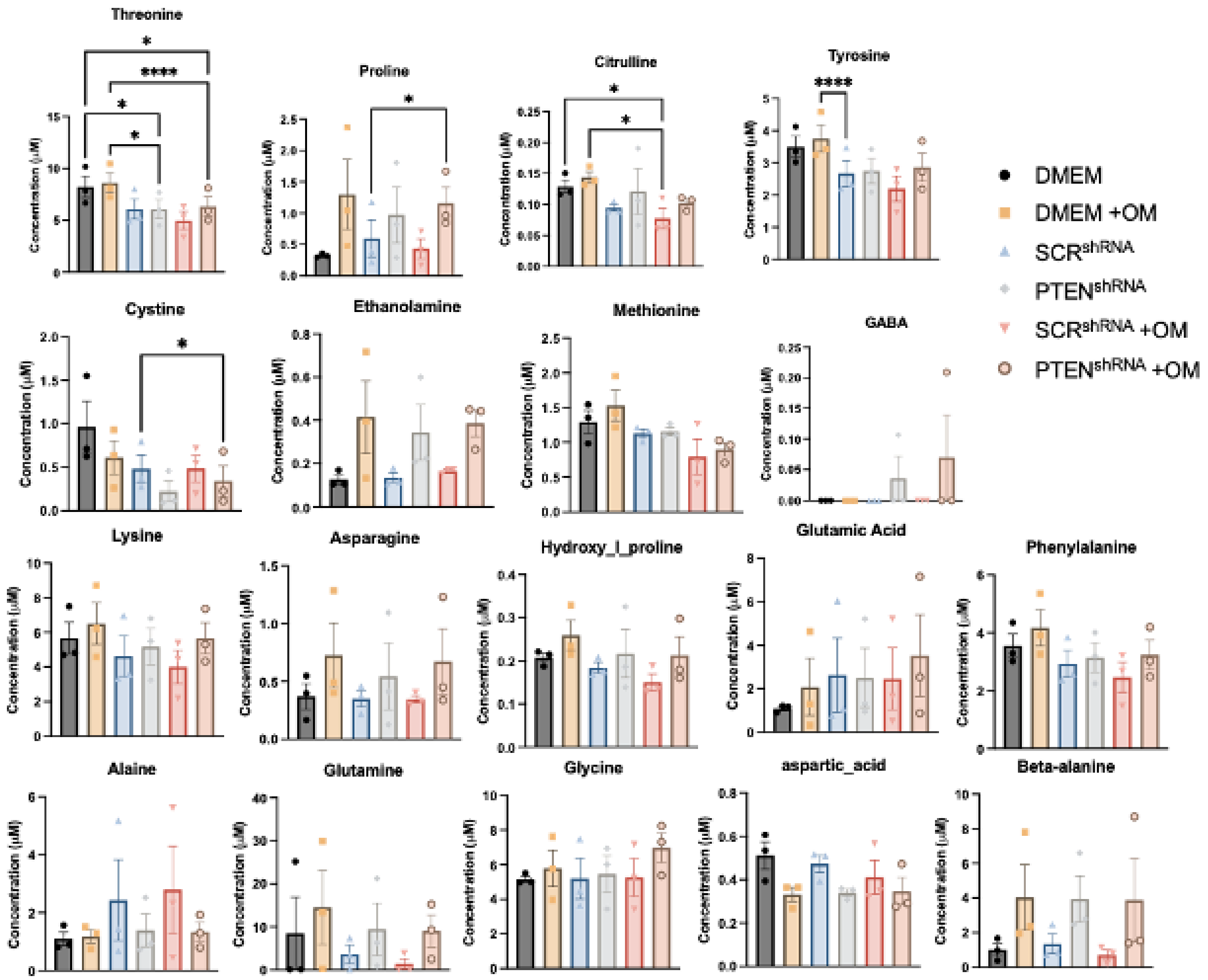
